# Transcranial direct current stimulation (tDCS) promotes myelin repair and plasticity in the mouse motor cortex

**DOI:** 10.64898/2026.01.20.700602

**Authors:** Enrica Boda, Federica Marchiotto, Alessandro Pigozzi, Annalisa Buffo, Mario Buffelli, Marco Cambiaghi

## Abstract

Cortical myelin loss and oligodendroglial dysfunction, hallmarks of numerous neurodegenerative and psychiatric disorders, compromise executive, sensory, and cognitive functions by altering circuit activity and integrity. Promoting cortical myelin repair therefore represents a critical therapeutic goal. In this frame, modulation of neuronal activity through non-invasive brain stimulation may represent a promising non-pharmacological approach to support remyelination.

To address this issue, we investigated the effects of anodal transcranial direct current stimulation (A-tDCS) in a mouse model of unilateral myelin injury in the motor cortex. A-tDCS was delivered over the contralateral uninjured cortex during a bilateral motor task (locomotion), to enhance the physiological activity of interhemispheric M1–M1 projections and indirectly stimulate the lesioned cortex. When applied during the late phase of demyelination and the onset of repair, A-tDCS accelerated remyelination and increased oligodendroglial survival and maturation in the lesioned cortex, compared with unstimulated controls. Increased myelin was also detected in the directly stimulated uninjured cortex despite unchanged oligodendroglial cell numbers, suggesting *de novo* myelin formation or remodeling of existing internodes. These structural changes were accompanied by preservation of interhemispheric M1–M1 functional connectivity upon injury.

Collectively, our findings demonstrate that A-tDCS promotes cortical myelin repair *in vivo*, supporting its therapeutic potential for disorders characterized by myelin damage. Notably, the observed effects of tDCS on myelin plasticity in the uninjured cortex – reported here for the first time - reveal a previously unrecognized mechanism underlying tDCS-mediated neuromodulation.

**Highlights:**

- Anodal tDCS (A-tDCS) promotes remyelination in the mouse motor cortex (M1) and restores M1–M1 synchrony after demyelinating injury
- A-tDCS enhances oligodendroglial survival and maturation in the injured motor cortex
- A-tDCS induces myelin plasticity in the uninjured cortex

## Introduction

In the nervous system, the speed of action potential conduction along axons is directly related to the extent of myelination (i.e. the length and thickness of the myelin sheaths – also called *internodes*; (Friede, 1986; Waxman, 1977). Central Nervous System (CNS) myelination is thus considered one of the major contributors to the evolutionary success of vertebrates, being uniquely expanded in the human brain and essential for the development and execution of motor, sensory, and cognitive functions (Tomassy et al., 2016). Moreover, myelin plasticity (i.e. addition of new myelin sheaths or remodeling of the existing ones in response to experience) plays a key role in the adaptation to environmental changes and different types of learning and memory (Eugenin von Bernhardi and Dimou, 2022; McKenzie et al., 2014; Pajevic et al., 2014; Pan et al., 2020; Shimizu et al., 2023). Yet, the neurosupportive functions of myelin and oligodendrocytes (OLs) - the myelinating cells of the CNS - go beyond just providing electrical insulation and include the regulation of neuronal excitability and survival *via* potassium buffering and metabolic supply (Schirmer et al., 2018; Simons et al., 2024).

Alterations in myelin deposition or myelin loss impair action potential conduction and compromise neuronal circuit integrity. Accordingly, disruption of myelin and OLs in cortical grey matter is a hallmark of several neurodegenerative and psychiatric disorders, such as Alzheimer’s Disease, Multiple Sclerosis (MS) and Major Depression (Boda, 2021; Depp et al., 2023; Lucchinetti et al., 2011; Magliozzi et al., 2023; Nave and Ehrenreich, 2014), and is associated with cognitive decline and disease progression (Stedehouder and Kushner, 2017; Timmler and Simons, 2019; Vucic et al., 2012). Spontaneous remyelination can occur following demyelination, driven by large by oligodendrocyte progenitor cells (OPCs) and partly by mature OLs (Franklin et al., 2021). After recruitment to demyelinated areas, OPCs amplify by active proliferation, differentiate into premyelinating and eventually mature OLs, engage naked axons, and form new myelin sheaths.

However, upon aging or in case of repeated or chronic demyelination, myelin repair is often inefficient or incomplete, leading to persistent axonal dysfunction and neurodegeneration (Franklin and ffrench-Constant, 2017). Boosting remyelination is thus now considered a key therapeutic goal in regenerative medicine. Experimental studies have provided insights into the cell-intrinsic and environmental mechanisms governing the different steps of remyelination. Although a few of these target mechanisms have already entered clinical trials (Caprariello and Adams, 2022; Riboni-Verri et al., 2025), no pharmacological remyelination approach has demonstrated clear clinical benefits so far and effective therapies to restore myelin remain unavailable.

In this context, preclinical studies have shown that the modulation of neuronal activity may be a promising non-pharmacological strategy. In the healthy CNS, enhancing neuronal activity positively impacted onto OPC proliferation, differentiation, and myelination *in vivo* (Gibson et al., 2014; Kondiles et al., 2023; Mitew et al., 2018; Piscopo et al., 2018). Consistently, in animal models of myelin injury, the optogenetic and chemogenetic activation of neurons promoted oligodendrogenesis and remyelination (Luo et al., 2021; Ortiz et al., 2019). Although the mechanisms underlying these effects are largely unknown, these findings pave the path for testing whether non-invasive manipulation of neuronal activity might be exploited to boost OL generation and myelin repair.

Long-term, non-invasive multisensory (visual and auditory) stimulation at 40 Hz (gamma frequency) has been shown to reduce myelin loss and promote oligodendrogenesis following toxic demyelination (Rodrigues-Amorim et al., 2024), highlighting the therapeutic potential of non-invasive brain stimulation (NIBS). In this frame, repetitive transcranial magnetic stimulation (rTMS) promoted OL survival and maturation and increased myelination (Cullen et al., 2019), while decreasing nodal length (Cullen et al., 2021) in the adult healthy mouse brain. rTMS also induced transcriptional changes in oligodendroglial and myelin-related genes *in vivo* (Ong and Tang, 2025) and *ex-vivo* (Weiler et al., 2023). Moreover, depending on the time of intervention, rTMS and repetitive trans-spinal magnetic stimulation (rTSMS) either prevented myelin/OL loss (Chen et al., 2025; Ramírez-Rodríguez et al., 2023; Sherafat et al., 2012; Sun et al., 2018; Yang et al., 2020) or promoted myelin repair (Fabres et al., 2025; Nguyen et al., 2024; Semprez et al., 2025; Z. Wang et al., 2021) in preclinical models of myelin injury.

The effects of other NIBS approaches – and particularly of transcranial direct current stimulation (tDCS) - on myelin damage and repair have been explored only marginally so far. tDCS is emerging as a promising tool for the treatment of various neurological and neuropsychiatric conditions (Cambiaghi et al., 2023). tDCS involves the application of a weak, transcranially delivered direct current, flowing from positive (anodal) to negative (cathodal) poles with neuromodulatory effects on cortical excitability (Peruzzotti-Jametti et al., 2013). Early evidence obtained in the primary motor cortex (M1) showed that anodal (A-tDCS) and cathodal tDCS (C-tDCS) had opposite effects, with anodal stimulation increasing and cathodal stimulation decreasing M1 excitability beyond the stimulation period in both humans (Nitsche and Paulus, 2000) and mice (Cambiaghi et al., 2010). In models of myelin injury, C-tDCS applied during acute or sub-acute stages enhanced myelin preservation and increased OPC recruitment (Braun et al., 2016; Marenna et al., 2022; Zhang et al., 2020). So far, the therapeutic potential of A-tDCS on myelin has been explored at preclinical level in specific contexts, i.e. together with mesenchymal stem cell transplantation (Mojaverrostami et al., 2022) or combined with intense physical activity along the visual pathway (Rossi et al., 2026, 2024). Overall, these studies showed a positive effect of A-tDCS on the functional outcomes of injury, although with heterogenous effects on myelin recovery. Of note, A-tDCS has been able to improve cognitive deficits in MS patients. Yet, the contribution of myelin preservation or repair in this effect is unknown (Charvet et al., 2025; Mattioli et al., 2016). Thus, there is a strong rationale to further investigate whether tDCS can be proposed as a strategy to boost remyelination and how it exerts its effects.

To address this issue, we investigated the effects of A-tDCS in a mouse model of unilateral demyelinating injury within the motor cortex. A-tDCS was applied over the contralateral (uninjured) cortex to positively modulate the physiological activity of interhemispheric M1-M1 fibers during a motor task (i.e. low-intensity locomotion) involving both hemispheres. This stimulation paradigm is known to modulate neural activity in the non-directly stimulated hemisphere via interhemispheric connections and therefore targets the lesioned cortex indirectly (Marchiotto et al., 2025). Results showed that, when applied during the late demyelination phase and in concomitance with myelin repair initiation, A-tDCS accelerated remyelination and increased oligodendroglia survival and maturation in the lesioned cortex, compared to unstimulated controls. Moreover, increased myelin was also observed in the directly stimulated uninjured cortex, despite unchanged oligodendroglial cell numbers, suggesting novel myelin formation and/or remodeling of the existing internodes. Such changes were accompanied by the restoration of M1-M1 functional connectivity.

Overall, these data demonstrate that A-tDCS promotes myelin repair and myelin plasticity in the mouse cortex, supporting its potential application in myelin disorders and unveiling a new target mechanism for the therapeutic and neuromodulatory actions of tDCS.

## Material and Methods

### Animals and experimental design

C57BL/6JOlaHsd male mice (12 weeks, 20–25 g at the start of the experiment; Envigo, San Pietro al Natisone, Italy) were used for this study. Mice were housed in the vivarium under standard conditions (12-hr light/12-hr dark cycle at 21°C) with food and water ad libitum. The project was designed according to the guidelines of the NIH, the European Communities Council (2010/63/EU) and the Italian Law for Care and Use of Experimental Animals (DL26/2014). It was also approved by the Italian Ministry of Health and the Bioethical Committee of the University of Turin. The study was conducted according to the ARRIVE guidelines. The experimental plan consisted of a lysolecithin (lysophosphatidylcholine, LPC, Sigma-Aldrich) -mediated myelin lesion and A-tDCS/sham stimulation combined with rotarod (see below). Animals were sacrificed at 7, 14 and 28 days post-injury (dpi) and transcardially perfused. To fate map cells generated during the post-injury phase, we employed the thymidine analog 5-bromo-2-deoxyuridine (BrdU, Sigma Aldrich) that is incorporated in the DNA during the S-phase of the cell cycle. BrdU (100 mg/kg body weight) was injected i.p. at 7 dpi – i.e. 2 hours (h) after the first session of stimulation (Fig. 1).

**Figure 1.**
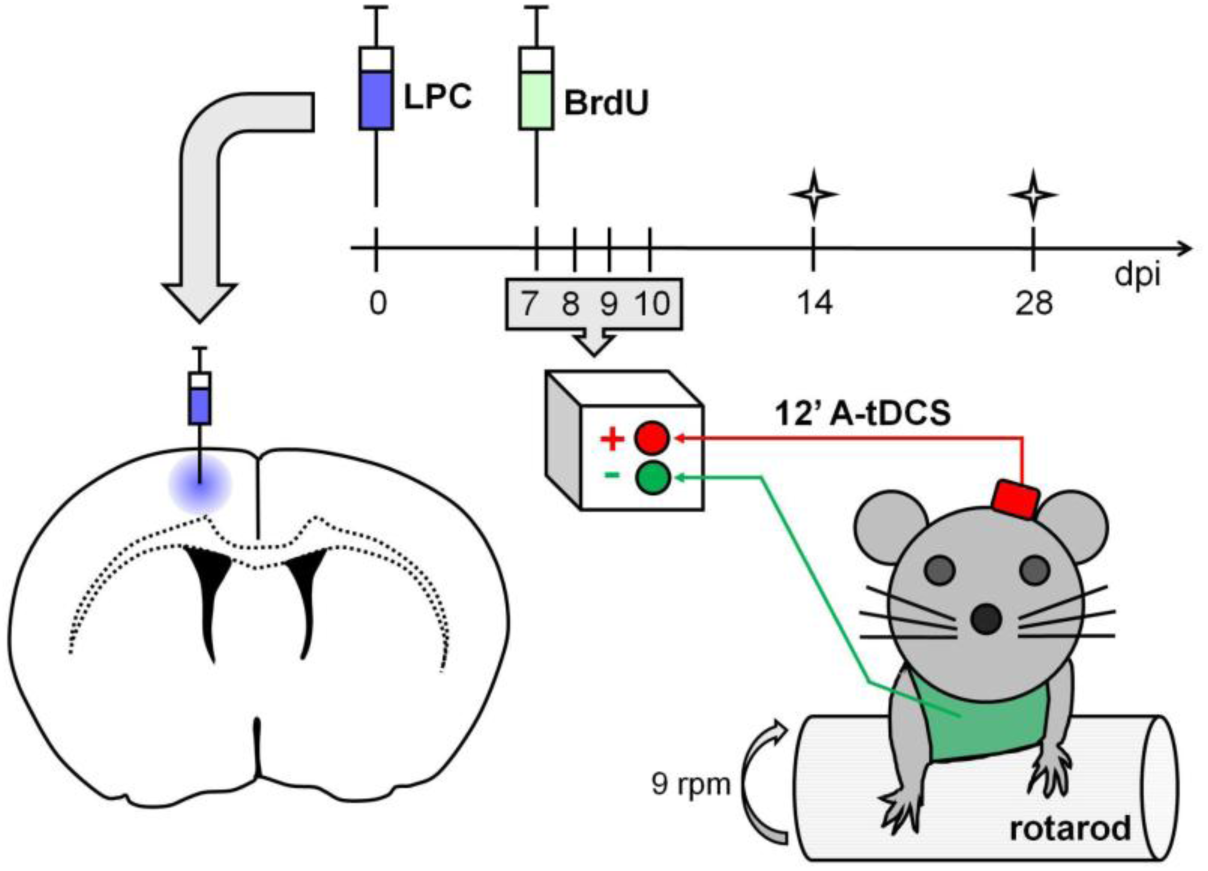
Experimental design: application of A-tDCS coupled with locomotion in a model of toxic demyelination. Contralateral A-tDCS combined with locomotion in mice subject to unilateral LPC-induced myelin injury in M1 cortical gray matter. To fate map cells generated during the stimulation phase, BrdU was injected i.p. at 7 dpi – i.e. 2 hours (h) after the first session of stimulation. The recordings of extracellular field potentials were made in awake mice at rest, at baseline (i.e. 7 dpi before the initiation of the stimulation) and at 28 dpi. Mice were sacrificed at 7, 14, and 28 dpi for histological analyses. Abbreviations: A-tDCS, anodal transcranial direct current stimulation; BrdU, 5-bromo-2-deoxyuridine; dpi, days post-injury; LPC, lysolecithin; M1, primary motor cortex.

### Acute LPC-mediated myelin injury and electrodes implant

Surgeries were carried out under general anesthesia obtained by i.p. administration of ketamine-xylazine (80 mg/kg and 5 mg/kg, respectively). Acute demyelination was obtained by a right unilateral stereotaxic microinjection of 1 μl of 1% LPC (Sigma-Aldrich) in 0.9% NaCl (saline) solution into the grey matter of the primary motor cortex (M1) at coordinates: 1 mm mediolateral, 0.1 mm rostral to bregma and at 0.8 mm from the cortical surface. tDCS electrode consists of a tubular plastic jack (inner area 4,5 mm2) that was dental cemented over the left M1 area, 2 mm lateral to Bregma (Cherchi et al., 2022). To record extracellular field potentials in awake mice, tungsten wires (50 μm Ø) were implanted in the right and left M1 cortex, according to the following coordinates: AP = +0.3 mm, ML = 1.0 mm from Bregma and DV = -0.5 mm from the brain surface (layer III), with the reference screw electrode over the cerebellum. To verify the correct location of the electrodes Cresyl Violet staining was performed *ex-post* in each mouse (see Fig. 7A,B).

### Transcranial direct current stimulation (tDCS) combined with locomotion

After mouse handling and habituation to the rotarod (2-6 days post injury, dpi), starting 7 days after LPC-mediated lesion, mice were subjected to anodal tDCS (A-tDCS) at a current intensity of 250 μA for 12 minutes for four consecutive days between 10 and 12 am. To deliver A-tDCS the electrode was filled with saline solution (0.9% NaCl) and the counter electrode (a saline-soaked sponge) was applied over the ventral thorax by using a rubber-made corset (Marchiotto et al., 2025). Animals were randomized to 1) receive A-tDCS combined with rotarod at constant speed (9 rpm) (LPC tDCS+walk), 2) receive sham tDCS (no current delivered) combined with rotarod (LPC walk) or 3) receive sham tDCS (no current delivered) without rotarod (LPC).

### Histological procedures

Animals were deeply anesthetized (ketamine-xylazine 90 mg/kg and 9 mg/kg, respectively) and transcardially perfused with 4% paraformaldehyde (PFA; Sigma-Aldrich) in 0.1 M phosphate buffer (PB). Brains were postfixed overnight, cryoprotected, and processed according to standard immunohistological procedures (Boda et al., 2022). Brains were cut in 30 μm thick coronal sections collected in PBS and then stained to detect the expression of different antigens: NG2 (rb, 1:200, Sigma-Aldrich AB5320); BCAS1 (rb, 1:500, Synaptic Systems); GST-pi (rb, 1:500, Eppendorf); MBP (m, Smi-99 clone, 1:1000, Biolegend); BrdU (m, 1:10,000, Sigma-Aldrich MAB3510); CASPR (rb, 1:500, Abcam ab34151). Incubation with primary antibodies was made overnight at 4 °C in PBS with 1-2% Triton-X 100. To allow BrdU recognition, slices were treated with citrate buffer pH 6.0 for 5 min at 90°C before adding primary antibodies. The sections were then exposed for 3 h at room temperature (RT) to secondary Cy3, Cy5 (Jackson ImmunoResearch Laboratories) or Alexa Fluor 488, Alexa Fluor 555 (Molecular Probes) -conjugated antibodies. 4,6-diamidino-2-phenylindole (DAPI, Fluka) was used to counterstain cell nuclei. After processing, sections were mounted on microscope slides with Tris-glycerol supplemented with 10% Mowiol (Calbiochem).

## Image acquisition and data analysis

Histological specimens were examined using a Nikon C1 confocal microscope (with the associated EZ-C1 Ver3.90 software, Nikon), or a Leica TCS SP5 confocal microscope (with the associated LAS AF 4.0 software, Leica Microsystems). Confocal images (1024 × 1024 pixels) were all acquired at 40×, with a speed of 100–200 Hz and 67.9 - 34.32 μm pinhole size.

Spectral confocal reflectance microscopy (SCoRe), which takes advantage of the high refractive index of myelinated tissue (Schain et al., 2014), was used to detect and quantify compact myelin using reflected light, as described in (Craig et al., 2024). All reflectance images of myelin were captured on a Leica TCS SP5 confocal microscope with an acoustic-optical beam splitter (AOBS) with a 70/30 reflectance/transmission (RT) ratio. The AOBS collected data from laser wavelength of 633 nm using one PMT photodetector with a detection bandwidth centered around the 633 nm laser wavelength.

Adobe Photoshop 6.0 (Adobe Systems, San Jose, CA) was used to assemble the final plates. Quantitative evaluations were performed on confocal images with Fiji/ImageJ (Research Service Branch, National Institutes of Health, Bethesda, MD; available at http://rsb.info.nih.gov/ij/). To analyze the expression level of MBP and of myelin SCoRE signal, the positive fractioned area (i.e. the percentage of positive pixels) was quantified in 40x confocal image stacks comprising 20 optical slices 0.99 µm thick (for MBP) or 10 optical slices 0.99 µm thick (for myelin SCoRE). Density of oligodendroglial (NG2+, BCAS1+, GSTpi+) cells was calculated as number of cells per mm2 in 40x confocal image stacks comprising 20 optical slices 0.99 µm thick. To analyze the percentages of coexpression of NG2/BrdU, BCAS1/BrdU and GSTpi/BrdU, all BrdU+ cells present in the lesion area were inspected. Lesion boundaries were assessed using the anti-MBP staining. To scan the entire rostro-caudal lesion extension, all sections including the lesion were considered for the quantifications. At least three animals and at least three sections per animal were analyzed for each experimental condition.

### LFP recordings and analysis

The recordings of extracellular field potentials were made in awake resting mice, at baseline (i.e. 7 dpi before the initiation of the stimulation) and at 28 dpi. A customized Python script was used for offline coherence analysis, for which three 2-seconds epochs were averaged. Differences in coherence were obtained by subtracting mean coherence values (post-stimulation – pre-stimulation) and measured within the following frequency bands: delta (1-4 Hz), low-theta (4-8 Hz), high-theta (8-12 Hz), alpha (12-20 Hz), and slow-gamma (20-30 Hz) (Cambiaghi et al., 2016; Nasini et al., 2023).

### Statistical Analysis

Statistical analyses were carried out with GraphPad Prism 9 (GraphPad software, Inc). The Shapiro-Wilk test was first applied to assess whether data followed a normal distribution. When normally distributed, unpaired Student’s t-test (to compare two groups) or One-way/Two-ways Anova (for multiple group comparisons) followed by Bonferroni’s post-hoc analysis, were used. Alternatively, when data were not normally distributed, the Mann–Whitney U-test was used. Statistics also included Chi-square test to compare frequencies. In all instances, P *<* 0.05 was considered as statistically significant. Histograms represent mean ± standard error (SE). Statistical differences were indicated with * P *<* 0.05, **P *<* 0.01, ***P *<* 0.001.

## Results

### tDCS accelerates myelin repair in a model of myelin injury in the cortical grey matter

To investigate whether tDCS affects myelin repair, we exploited a mouse model of unilateral LPC-induced myelin lesion within the primary motor cortex (M1), where acute demyelination is followed by spontaneous remyelination sustained by resident OPCs and spared OLs. Since the LPC injury model was most commonly applied to white matter regions (Plemel et al., 2018), we first investigated whether this lipid-disrupting agent similarly preserves axons in M1 grey matter. By quantifying the immunolabeling obtained with the pan-axonal antibody marker SMI312, which is directed against highly phosphorylated axonal neurofilaments (Lee et al., 1987), we detected a sizable axonal sparing after LPC injury (about 85% at 28 dpi; Suppl. Fig. 1A, B), thus validating the suitability of this model for studying myelin repair also in the cortical grey matter.

Mice received A-tDCS over the contralateral (uninjured) cortex in combination with low-intensity physical activity (i.e. walking on a rotarod) daily from 7 to 10 dpi, a time window corresponding to the late demyelination phase and the initiation of reparative processes. This stimulation protocol allows an indirect modulation of the lesioned region *via* interhemispheric connections (Marchiotto et al., 2025). To increase the probability to detect even fine differences in lesion repair, the effects of this manipulation were studied at 14 dpi - when we observed that oligodendroglia maturation was still ongoing and remyelination had not yet occurred (Fig. 2B,C,E,F) – and at 28 dpi – when myelin repair had not reached a plateau (Fig. 2B,C,E,F). Consistent with previous findings (Zoupi et al., 2021), in the cortical grey matter LPC injection did not result in an obvious myelin loss before 14 dpi, when a 40% decrease of the myelin basic protein (MBP) immunolabeling was observed in both supragranular (I-III) and infragranular (V-VI) layers of the lesioned cortex, compared to the intact ones (Fig. 2A-C). Consistently, the reflection signal of myelinated axons appeared reduced by 50% in both superficial and deep cortical layers, as assessed by SCoRe microscopy (Fig. 2D-F). Of note, post-injury application of A-tDCS combined with physical activity (tDCS+walk) – but not physical activity *per se* (i.e. walk; Suppl. Fig. 2A) – accelerated the restoration of MBP expression (Fig. 2A-C). and myelin deposition (Fig. 2D-F) in both supragranular and infragranular layers at 28 dpi, compared to controls. These data indicate that application of A-tDCS in the post-acute phase promotes myelin repair in the cortical grey matter.

**Supplementary Figure 1.**
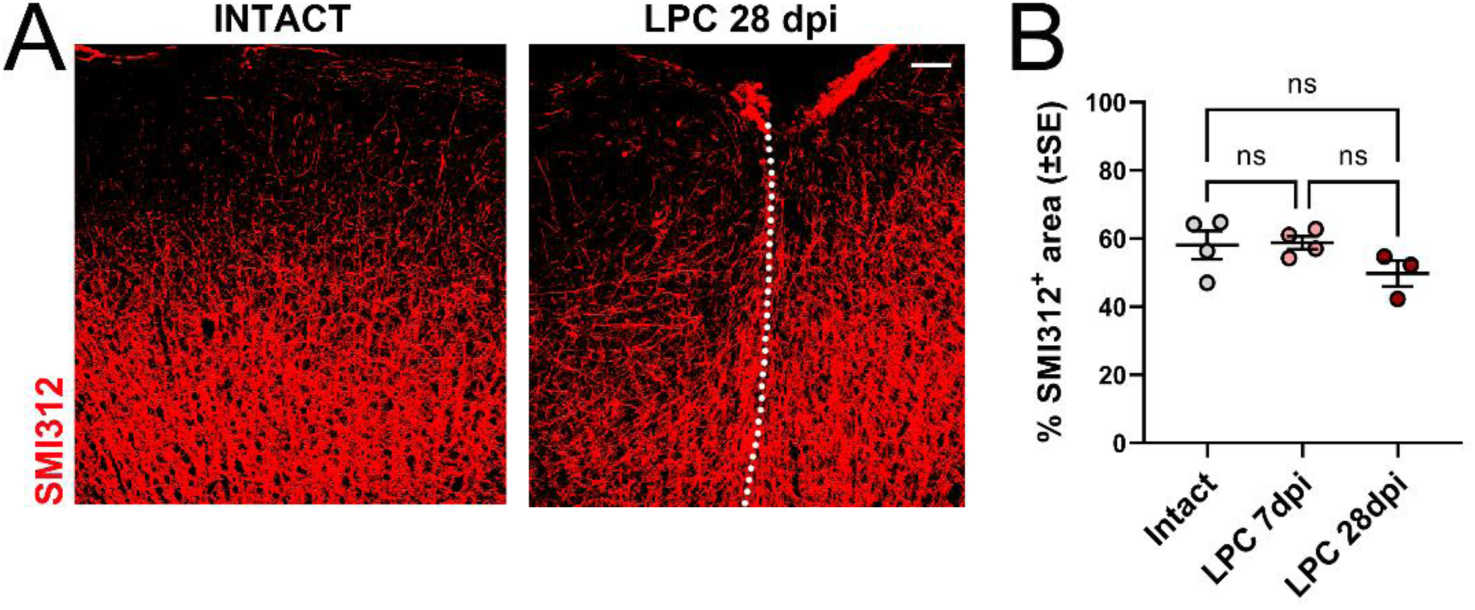
Axon sparing following LPC-induced myelin injury in M1 grey matter. (A) Representative images of SMI312 (red) expression pattern in the intact (first panel) and lesioned (28 dpi, second panel) M1 cortex. The dotted line indicates the injection track. Scale bar: 50 µm. (B) Quantification of the percentage of SMI312+ area in the intact *vs*. injured cortex at 7 and 28 dpi. Quantification includes both supragranular (i.e. I-III) and infragranular (i.e. V-VI) layers. Each dot represents an individual mouse. Differences were assessed by One-way Anova followed by Bonferroni’s Multiple Comparison Test.

**Figure 2.**
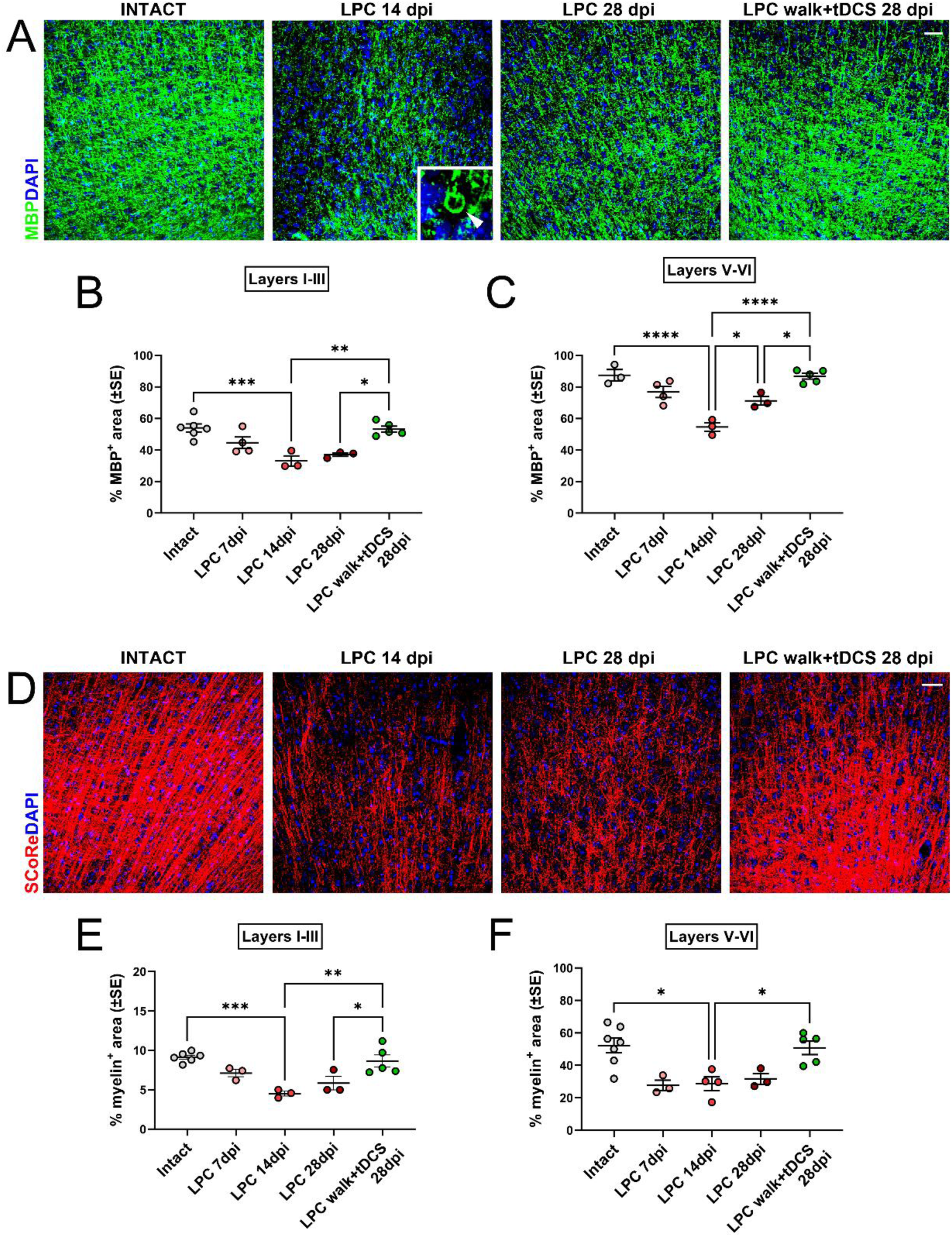
A-tDCS accelerates myelin recovery following myelin injury. (A) Representative images of MBP (green) expression pattern in the intact (first panel) and lesioned M1 cortex. Note the strong decrease of MBP+ signal as well as the presence of MBP+ debris and degenerative “ovoidal figures” (inset, indicated by arrowhead) in unstimulated mice at 14 dpi (2nd panel). At 28 dpi, MBP+ labeling increased in mice exposed to tDCS+walk (LPC walk+tDCS) compared to unstimulated lesioned mice (LPC). (B,C) Quantification of the percentage of MBP+ area in supragranular (i.e. I-III, B) and infragranular (i.e. V-VI, C) layers of the lesioned cortex. (D) Representative images of the SCoRe signal (i.e. compact myelin) in the intact (first panel) and lesioned M1 cortex at 14 and 28 dpl. At 28 dpi, the SCoRe signal increased in mice exposed to tDCS+walk (LPC walk+tDCS) compared to unstimulated lesioned mice (LPC). DAPI (blue) counterstains cell nuclei in A and D. Scale bars: 50 µm. (E,F) Quantification of the percentage of SCoRe+ area in supragranular (i.e. I-III, E) and infragranular (i.e. V-VI, F) layers of the lesioned cortex. Each dot in (B,C,E,F) represents an individual mouse. Differences were assessed by One-way Anova followed by Bonferroni’s Multiple Comparison Test. *P < 0.05; **P < 0.01; ***P < 0.001; ****P < 0.0001.

### tDCS sustains post-injury oligodendroglia survival and maturation

To assess whether the observed effect of tDCS on myelin repair resulted from the modulation of the dynamics of oligodendroglia proliferation and maturation, and/or from the prevention of oligodendroglia loss during the course of de-/re-myelination, we analyzed the densities of cells expressing markers of immature OPCs (i.e. NG2), premyelinating (i.e. BCAS1) and mature (i.e. GSTπ) OLs at the lesion site at 14 and 28 dpi. NG2-positive (+) and GSTπ+ cells were counted in both supragranular and infragranular layers, whereas the intricate labeling of myelinated axons allowed a reliable assessment of BCAS1+ cell bodies only in layers I-III.

Consistent with myelin/MBP data reported above, at 14 dpi unstimulated mice showed an ongoing and incomplete regenerative response to injury, with NG2+ OPCs and mature GSTπ+ OLs not differing from those in the intact cortex and intermediate phenotypes (i.e. premyelinating BCAS1+ cells) displaying a 6-fold increase (Fig. 3B,C,E,H,G). Of note, at this post-injury stage, the application of tDCS+walk resulted in a global expansion of the entire oligodendroglial lineage. Specifically, in both supragranular and infragranular layers NG2+ OPC density showed a 45% increase in tDCS+walk -exposed mice, compared to unstimulated mice (Fig. 3A-C). tDCS+walk displayed an even more pronounced effect on the density of premyelinating BCAS1+ (+65% in the supragranular layers; Fig. 3D,E) and mature GSTπ+ (a 2-fold increase in the supragranular and +40% in the infragranular layers) OLs, compared to controls (Fig. 3F-H). Yet, a general reduction in the abundance of all these oligodendroglia cell types was detected at 28 dpi in both unstimulated and, to a larger extent, tDCS+walk -exposed mice, suggesting the loss of supernumerary cells at later time points. Still, in tDCS+walk -exposed mice NG2+, BCAS1+ and GSTπ+ cells maintained a trend to be more abundant than in controls (Fig. 3B,C,E,H,G). Also in this case, physical activity *per se* (i.e. walk) did not show any significant effect on the density of oligodendroglia cells at lesion site (Suppl. Fig.2B-D).

**Figure 3.**
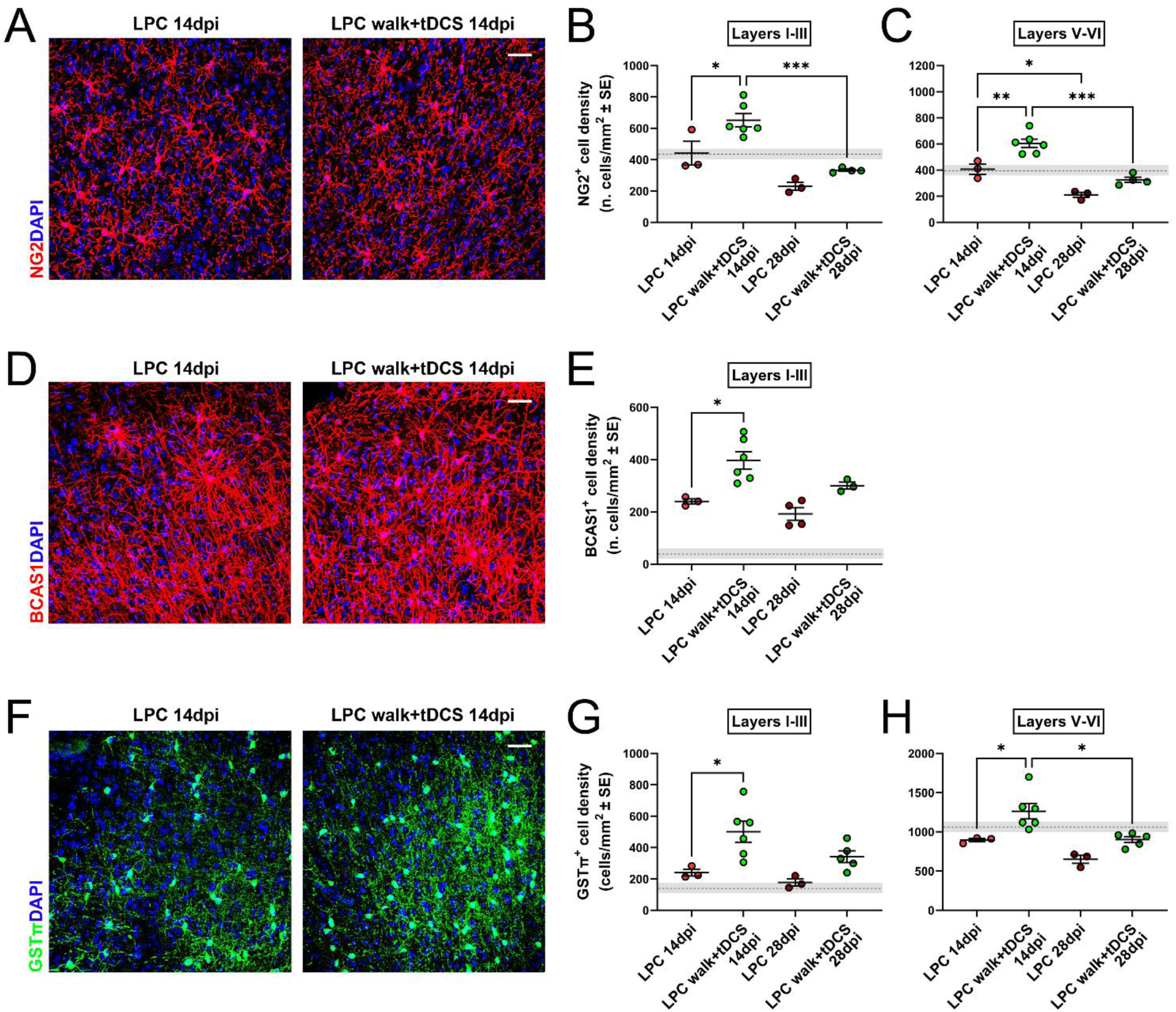
Post-injury application of A-tDCS coupled with physical activity results in the expansion of the entire oligodendroglial lineage. (A) Representative images of NG2+ (red) OPCs in the lesioned M1 cortex of unstimulated (LPC, first panel) and stimulated (LPC walk+tDCS, 2nd panel) mice at 14 dpi. (B,C) Quantification of the density of NG2+ OPCs in the supragranular (i.e. I-III, B) and infragranular (i.e. V-VI, C) layers of the lesioned cortex at 14 dpi and 28 dpi. (D) Representative images of BCAS1+ (red) premyelinating OLs in the supragranular layers of the lesioned M1 cortex of unstimulated (LPC, first panel) and stimulated (LPC walk+tDCS, 2nd panel) mice at 14 dpi. (E) Quantification of the density of BCAS1+ cells in the supragranular (i.e. I-III) layers of the lesioned cortex at 14 dpi and 28 dpi. (F) Representative images of GSTπ+ (red) mature OLs in the lesioned M1 cortex of unstimulated (LPC, first panel) and stimulated (LPC walk+tDCS, 2nd panel) mice at 14 dpi. (G,H) Quantification of the density of GSTπ+ mature OLs in the supragranular (i.e. I-III, G) and infragranular (i.e. V-VI, H) layers of the lesioned cortex at 14 dpi and 28 dpi. DAPI (blue) counterstains cell nuclei in (A,D,F). Scale bars: 50 µm. In (B,C,E,G,H), the dotted line and nearby grey area represent the mean cell density ± SE of the intact cortex, respectively. Each dot in (B,C,E,G,H) represents an individual mouse. Differences were assessed by Two-ways Anova followed by Bonferroni’s Multiple Comparison Test. *P < 0.05; **P < 0.01; ***P < 0.001; ****P < 0.0001.

**Supplementary Figure 2.**
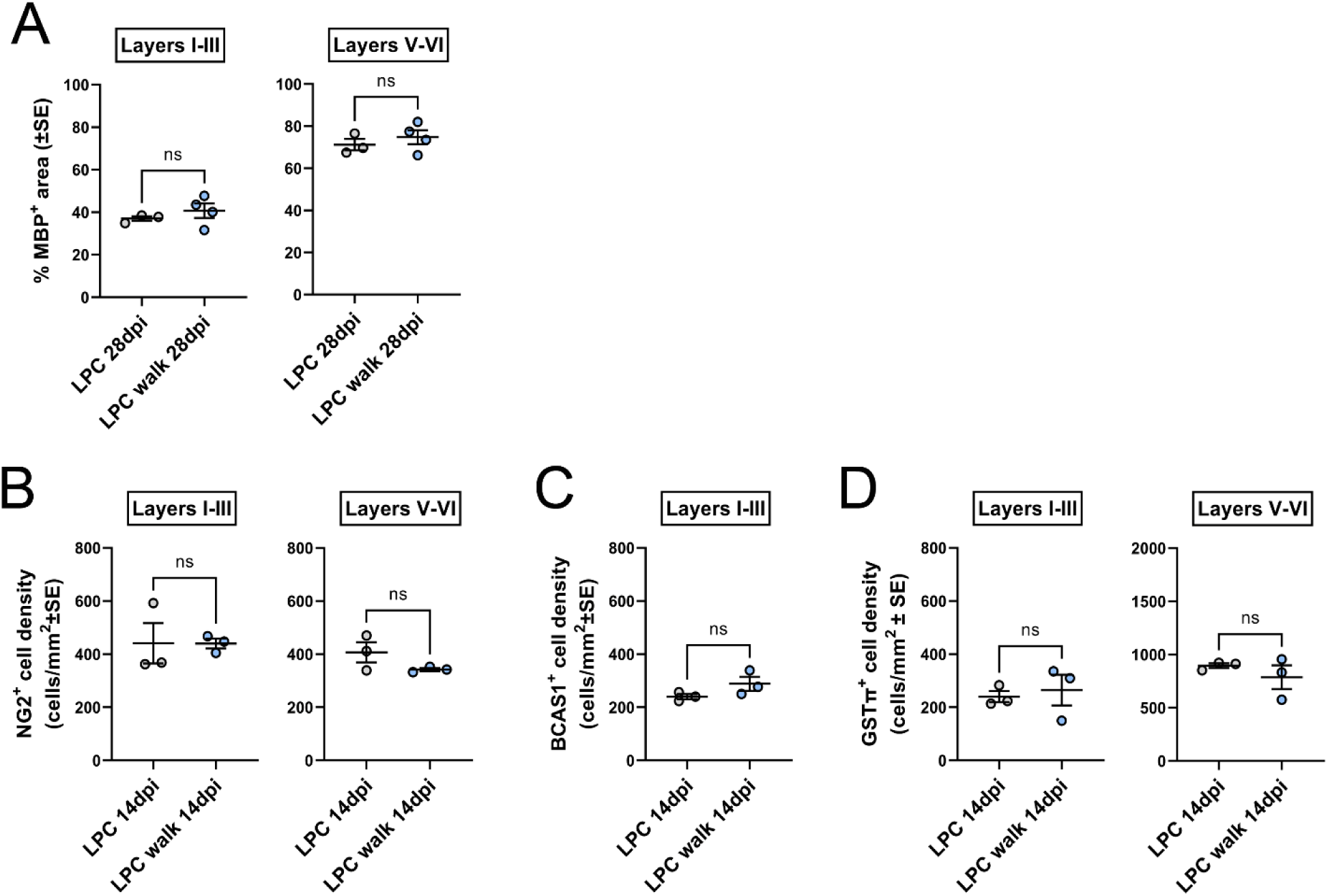
Physical activity *per se* does not accelerate myelin recovery following myelin injury. (A) Quantification of the percentage of MBP+ area in supragranular (i.e. I-III) and infragranular (i.e. V-VI) layers of the lesioned cortex of unstimulated (LPC) and mice subjected to sham stimulation+rotarod (LPC walk) at 28dpi. (B-D) Quantification of the density of NG2+ OPCs (B), BCAS1+ premyelinating OLs (C) and GSTπ+ mature OLs in the supragranular (i.e. I-III) and infragranular (i.e. V-VI) layers of the lesioned cortex of unstimulated (LPC) and mice subjected to sham stimulation+rotarod (LPC walk) at 14dpi. Each dot in (B,C,E,G,H) represents an individual mouse. Differences were assessed by unpaired Student’s t-test or Mann–Whitney U-test. *P < 0.05; **P < 0.01; ***P < 0.001; ****P < 0.0001.

The global expansion of the entire oligodendroglial lineage – including both immature and mature phenotypes – detected early after the stimulation window suggested a possible effect of tDCS on post-injury OPC proliferation. Yet, at 14dpi the density of cells born on the first day of tDCS+walk application (i.e. labeled by BrdU at 7 dpi) at lesion was not different in the tDCS+walk group *vs.* unstimulated controls (Fig. 4D). This clashes with the idea of a positive effect of tDCS+walk on OPC expansion during the second week after injury and instead supports a role for the intervention in promoting OPC survival during the course of de-/re-myelination.

**Figure 4.**
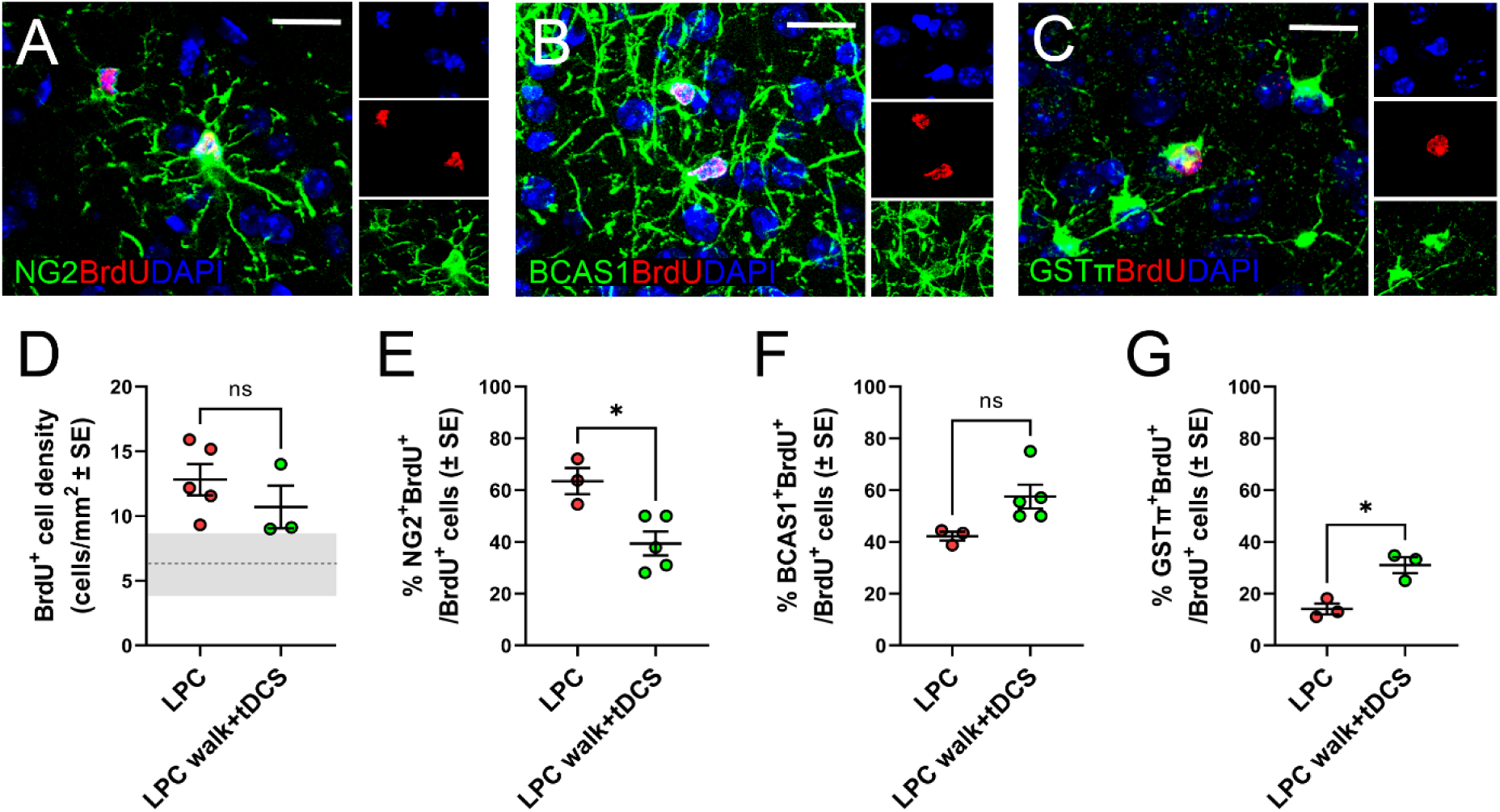
Accelerated post-injury maturation of oligodendroglia upon A-tDCS coupled with physical activity. (A-C) Representative image of NG2 (green in A), BCAS1 (green in B) and GSTpi (green in C) expression in BrdU+ (red) cells in LPC+walk+tDCS mice at 14 dpi. DAPI (blue) counterstains cell nuclei. Scale bars: 25 µm. (D) Quantification of the density of newborn (i.e. having incorporated BrdU at 7dpi) cells at lesion site at 14 dpi. The dotted line and nearby grey area represent the mean cell density ± SE of the contralateral uninjured cortex, respectively. (E,F) Quantification of the percentage of NG2+ OPCs and BCAS1+ premyelinating OLs among newborn (BrdU+) cells at 14 dpi. (G) Quantification of the percentage of GSTpi+ mature OLs among newborn (BrdU+) cells at 28 dpi. Each dot in (D-G) represents an individual mouse. Differences were assessed by unpaired Student’s t-test in (D) and by Chi-square test in (E-G). *P < 0.05; **P < 0.01; ***P < 0.001; ****P < 0.0001.

Increased numbers of premyelinating (i.e. BCAS1) and mature (i.e. GSTπ) OLs at lesion sites may also result from an accelerated maturation of cells generated during the reparative phases. Consistent with this idea, at 14 dpi, in the tDCS+walk group the fraction of BrdU+ cells that maintained NG2 expression was significantly lower (about 40%) compared to that of the controls (about 60%; Fig. 4A,E). In a complementary way, tDCS+walk promoted BrdU+ cell transition toward the premyelinating BCAS1+ stage (about 60% tDCS+walk *vs.* 40% ctrl; Fig. 4B,F), although almost none (i.e. ∼1%) of BrdU+ cells reached a mature GSTπ+ stage at 14 dpi (3 GSTπ+ cells over 204 BrdU+ cells in unstimulated mice; 1 GSTπ+ cell over 113 BrdU+ cells in tDCS+walk mice). Yet, at 28 dpi about 30% BrdU+ cells displayed GSTπ expression in the tDCS+walk group, with a 2-fold increase compared to the fraction of the controls (Fig. 4C,G). Taken together, these data indicate that post-acute application of tDCS sustains oligodendroglia survival and progression toward mature phenotypes during the reparative process.

### tDCS promotes myelin plasticity in the uninjured motor cortex

Activity-dependent changes in the cortical myelination pattern have been described in non-pathological conditions in both animal models and humans (Xin and Chan, 2020). Although myelin plasticity has been proposed as a possible substrate of tDCS effects (Antonenko et al., 2023), whether tDCS may actually affect myelin in the healthy brain has never been investigated so far. To address this issue, we examined MBP expression and myelin-associated reflection signal in the supragranular and infragranular layers of the directly-stimulated uninjured motor cortex (i.e. in the hemisphere contralateral to the lesion) at 14 dpi (i.e. 7 days after the initiation of tDCS+walk stimulation). While the highly-myelinated infragranular layers did not show any significant change, the most superficial and less myelinated layers displayed a significant increase in MBP labeling in the tDCS+walk group compared to unstimulated controls (Fig. 5A,B). Consistent results have been observed by myelin SCoRe imaging (Fig. 5C,D). Yet, at difference with what was observed in the lesioned cortex, such changes in the myelination pattern were not accompanied by any increase of immature NG2+ OPCs (Fig. 6A) or mature GSTπ+ OLs (Fig. 6B), suggesting novel myelin formation and/or remodeling of the existing internodes produced by a stable oligodendroglia population. Consistently, the density of newly-generated (i.e. BrdU+) cells did not change in tDCS+walk - exposed mice, compared to controls (Fig. 6C). Yet, in mice exposed to tDCS+walk, NG2+ cells displayed a hypertrophic arborization of processes (quantified as percentage of NG2+ area; Fig. 6D,E), indicating they set up a response to tDCS-induced increased activity in the local circuitries. Finally, in line with previous findings, physical activity *per se* (i.e. walk) did not result in myelin (i.e. MBP+ area; Suppl. Fig.3A) or OPC changes (i.e. NG2+ area; Suppl. Fig.3B).

**Figure 5.**
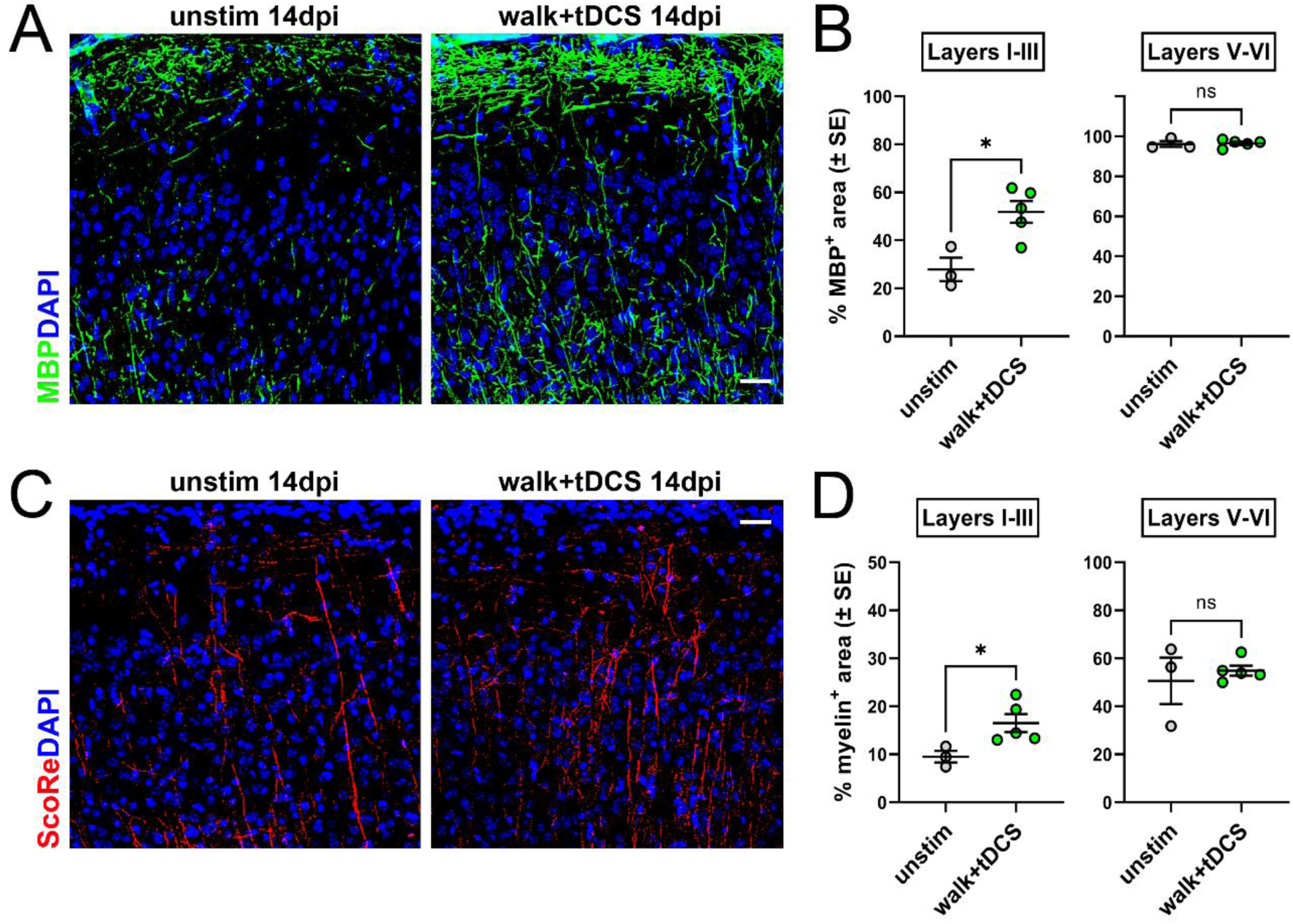
A-tDCS promotes myelin plasticity in the uninjured motor cortex. (A) Representative images of MBP (green) expression pattern in the unstimulated (unstim, first panel) and stimulated (walk+tDCS; 2nd panel) M1 cortex at 14 dpi (i.e. 7 days after the start of the stimulation period). Note the increase of MBP+ signal in stimulated mice (2nd panel). (B) Quantification of the percentage of MBP+ area in supragranular (i.e. I-III) and infragranular (i.e. V-VI) layers of the cortex. (C) Representative images of the SCoRe signal (i.e. compact myelin) in the in the unstimulated (unstim, first panel) and stimulated (walk+tDCS; 2nd panel) M1 cortex at 14 dpi (i.e. 7 days after the start of the stimulation period). DAPI (blue) counterstains cell nuclei in A and C. Scale bars: 50 µm. (D) Quantification of the percentage of SCoRe+ area in supragranular (i.e. I-III) and infragranular (i.e. V-VI) layers of the cortex. Each dot in (B,D) represents an individual mouse. Differences were assessed by unpaired Student’s t-test. *P < 0.05.

**Figure 6.**
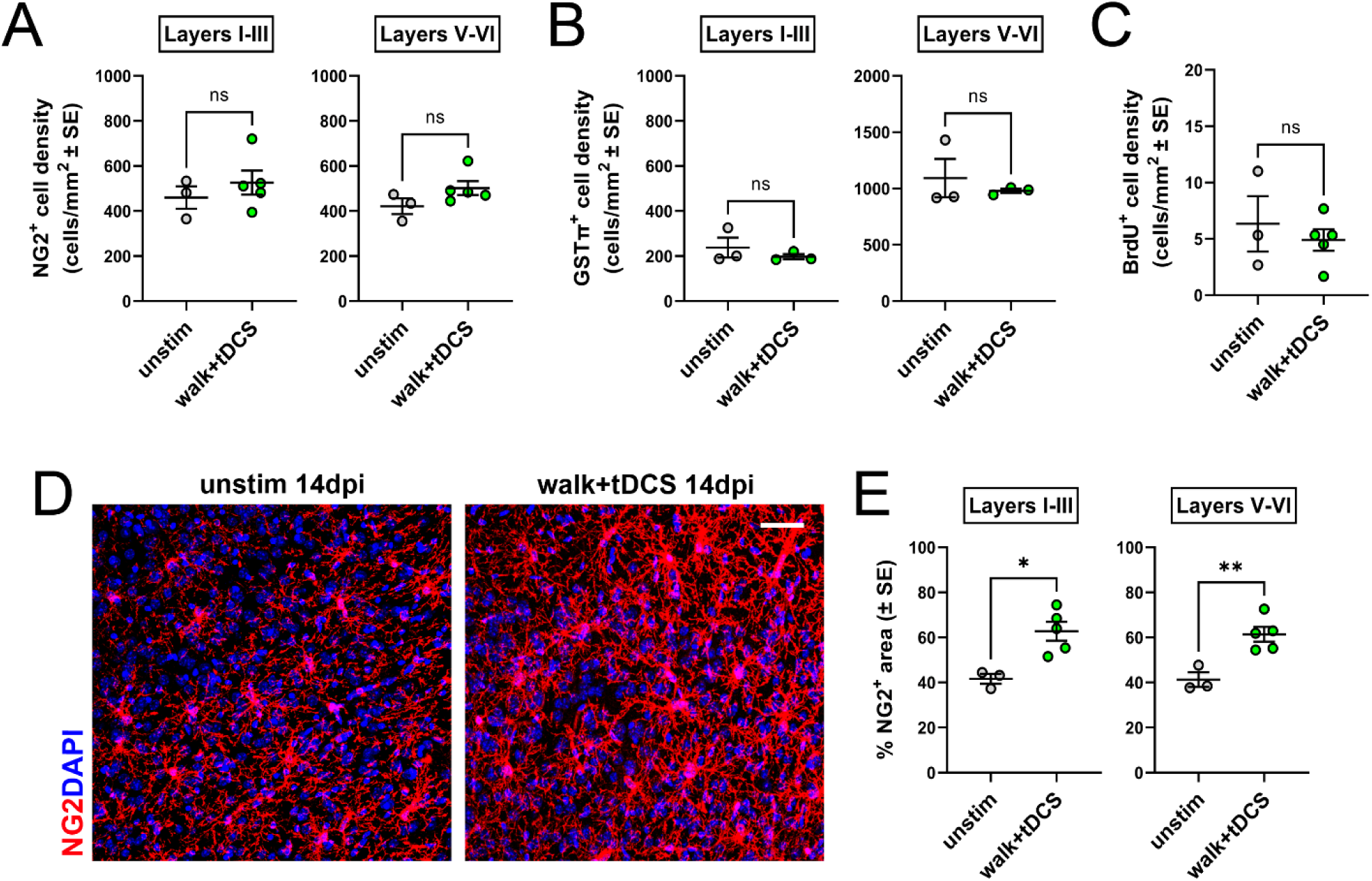
A-tDCS induces OPC hypertrophy in the uninjured motor cortex. (A-C) Quantification of the density of NG2+ OPCs (A), GSTπ+ mature OLs (B) and newborn BrdU+ cells (C) in the supragranular (i.e. I-III) and infragranular (i.e. V-VI) layers of the unstimulated (unstim) and stimulated (walk+tDCS) M1 cortex at 14 dpi (i.e. 7 days after the start of the stimulation period). (D) Representative images of NG2 (red) expression pattern in the unstimulated (unstim, first panel) and stimulated (walk+tDCS; 2nd panel) M1 cortex. Note the increase of NG2+ OPC processes in stimulated mice (2nd panel). DAPI (blue) counterstains cell nuclei. Scale bars: 50 µm. (E) Quantification of the percentage of NG2+ area in supragranular (i.e. I-III) and infragranular (i.e. V-VI) layers of the cortex. Each dot represents an individual mouse. Differences were assessed by unpaired Student’s t-test.

**Supplementary Figure 3.**
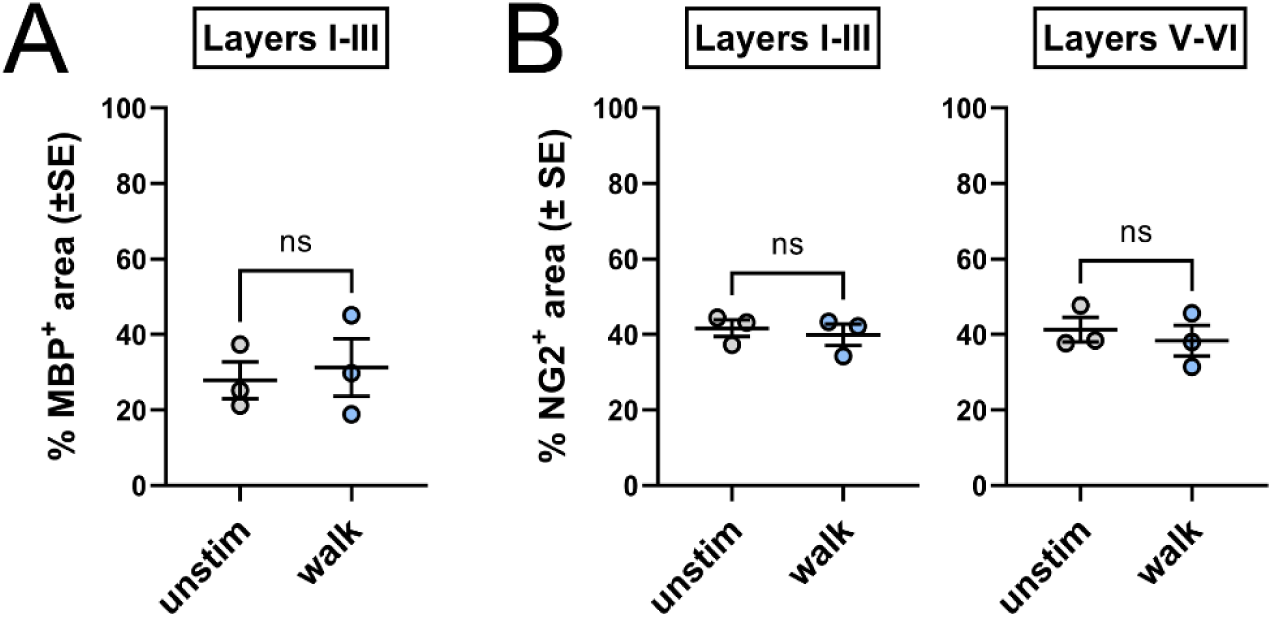
Physical activity *per se* does not promote myelin or OPC changes in the uninjured motor cortex. (A) Quantification of the percentage of MBP+ area in supragranular (i.e. I-III) layers of the cortex of unstimulated (unstim) and mice subjected to sham stimulation+rotarod (walk) at 14 dpi (i.e. 7 days after the start of the stimulation period). (B) Quantification of the percentage of NG2+ area in supragranular (i.e. I-III) and infragranular (i.e. V-VI) layers of the cortex of unstimulated (unstim) and mice subjected to sham stimulation+rotarod (walk) at 14 dpi. Each dot represents an individual mouse. Differences were assessed by unpaired Student’s t-test.

**Figure 7.**
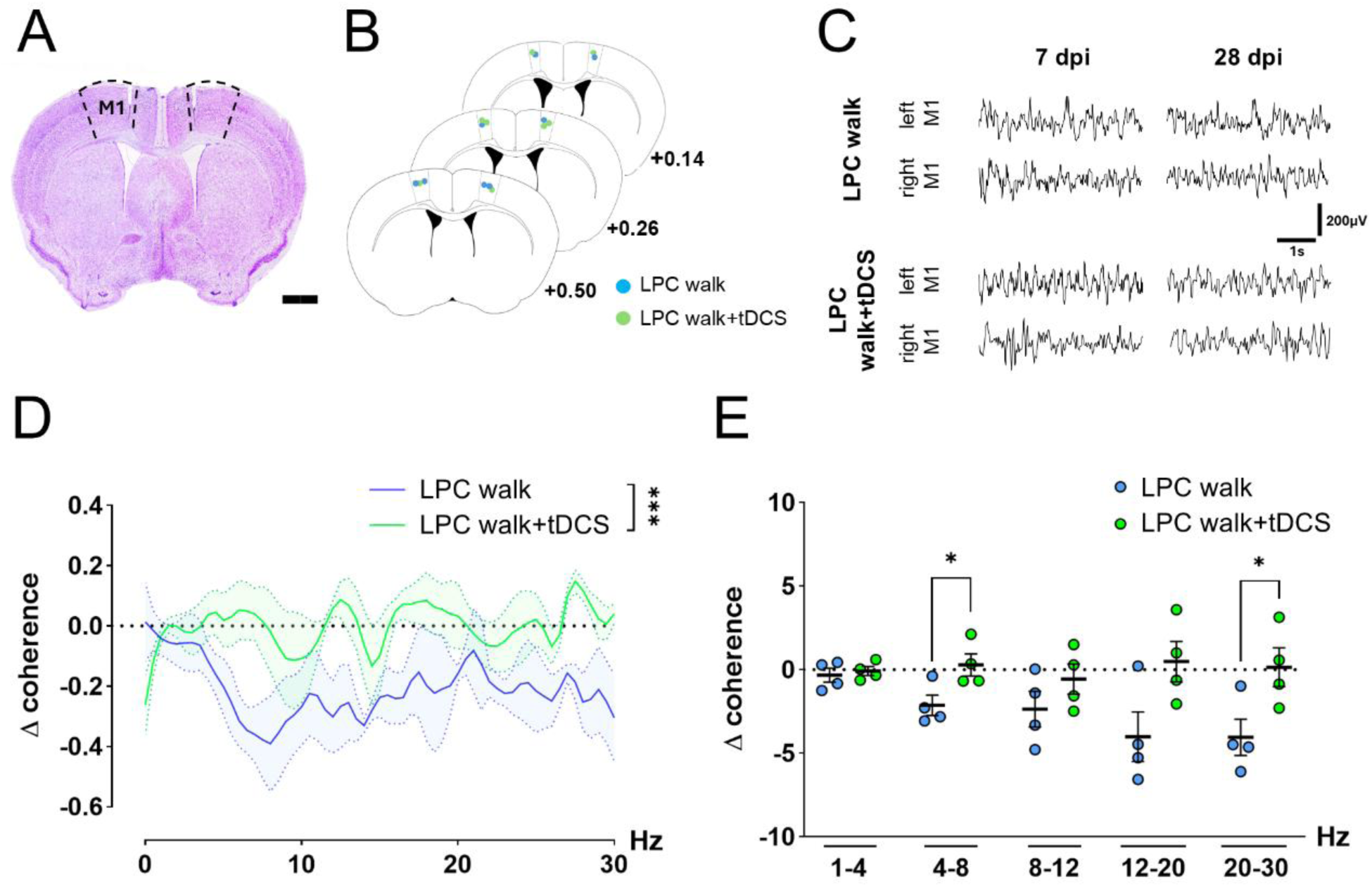
M1-M1 impaired interhemispheric synchrony is partially recovered by A-tDCS combined with physical activity. (A,B) Representative image of LFP recording electrodes in both M1 cortices (A) and location of electrode tips in the 2 groups (B). Scale bar in A: 1 mm. (C) Example of baseline and post-stimulation LFP traces recorded in the LPC-walk (top) and LPC-walk+tDCS (bottom) groups in the right/left M1. (D) After LPC lesion, the coherence difference between the post and the pre-tDCS period (Δ Coherence, 2-Ways-Anova) was recovered only in the LPC walk+tDCS group, with a significant increase in the 4-8 Hz and 20-30 Hz frequency ranges (E, unpaired Student’s t-test). *P < 0.05; ***P < 0.001.

Overall, these data show that A-tDCS promotes myelin plasticity and OPC morphological changes in the uninjured motor cortex.

### Post-injury application of tDCS impacts M1-M1 functional connectivity

As myelin regulation supports cortical synchronization, facilitating axonal conduction in long-range axons (Kato et al., 2020), we reasoned to investigate whether tDCS effect on myelin repair and plasticity were reflected by a functional correlate. Specifically, we recorded extracellular field potentials from the lesioned and the contralateral directly-stimulated M1 cortices to measure intercortical synchrony (Fig. 7A-C). Higher coherence (i.e. synchronization) between two regions is interpreted as a sign for functional coupling, while interhemispheric desynchronization is a valuable indicator of mono- or bi-lateral alterations (Marchiotto et al., 2025; Palmer et al., 2019). To address this issue, recordings were performed immediately before the application of tDCS (7 dpi, baseline) and at the latest time-point (28 dpi) in LPC walk+tDCS *vs* LPC walk mice. The difference in M1-M1 coherence between these observation points (Δ coherence) was used to evaluate the long-term effects of the 4-days A-tDCS treatment, in association with the observed effects on oligodendrocyte and myelin. In the LPC walk group, coherence was diminished at 28 dpi if compared to 7 dpi (Fig. 7D) in particular in the low-theta (4–8 Hz) and the slow-gamma (20–30 Hz) bands, which instead were preserved in the stimulated mice (Fig. 7E).

Together with the structural data, this outcome suggests that tDCS restores the functional impairment induced by LPC-induced demyelination.

## Discussion

Our study demonstrates that the combination of multisession (4-days) A-tDCS with moderate physical activity enhances myelin repair following focal demyelination and promotes myelin plasticity in the mouse motor cortex, resulting in a partial recovery of cortico-cortical interhemispheric synchrony. While former studies have shown a neuroprotective action of C-tDCS on myelin integrity (Braun et al., 2016; Marenna et al., 2022) and a positive effect of A-tDCS on myelin repair in the optic nerve and in the thalamic lateral geniculate nucleus (Rossi et al., 2026, 2024), to the best of our knowledge, this is the first demonstration that tDCS can influence (re-)myelination in the injured and healthy cerebral cortex.

Extensive myelin loss in the cortical gray matter is a common feature of MS and other neurological disorders (i.e. stroke, Alzheimer’s Disease, Major Depression), frequently exceeding the more extensively studied white matter demyelination (Boda, 2021; Depp et al., 2023; Magliozzi et al., 2023). Because a high load of cortical myelin lesions is associated with physical and cognitive decline (Stedehouder and Kushner, 2017; Timmler and Simons, 2019; Vucic et al., 2012), enhancing cortical remyelination represents a critical therapeutic goal. To examine whether tDCS influences myelin repair in the cortical grey matter, we injected LPC to induce a lipid disrupting myelinopathy within M1 (Plemel et al., 2018). Although the LPC lesion paradigm has been most frequently applied to white matter, previous studies have also used this model to induce focal demyelination in various cortical regions (Bonnefil et al., 2019; Cerina et al., 2017; Narayanan et al., 2018; Zoupi et al., 2021). Consistent with these reports, we found that, despite substantial preservation of axonal integrity - at least within the first month post-lesion - myelin loss and repair in the grey matter proceeded more slowly, and remyelination was markedly less efficient than in the white matter, with a minimal spontaneous recovery of MBP expression and no significant change in myelin ScoRe signal at 28 dpi. While this likely reflects a combination of oligodendroglial cell–intrinsic and environmental constrains shaping the grey matter response to myelin injury (Sherafat et al., 2021; Werkman et al., 2021), this tissue still retains a reparative potential that can be enhanced by stimulation.

Specifically, in our study, A-tDCS was applied over the contralateral (uninjured) cortex during locomotion, that - depending on the coordinated movements of both right and left limbs - requires bilateral motor cortex activation. In humans, unilateral tDCS over M1 can modulate the activity of the contralateral motor cortex (Lang et al., 2004). Moreover, building on evidence that neuronal networks co-activated by electrical stimulation exhibit heightened sensitivity to A-tDCS (Fritsch et al., 2010; Gellner et al., 2020; B. Wang et al., 2021), we have recently shown that combining unilateral A-tDCS with locomotion induces a bilateral increase of neuronal activity and structural plasticity (i.e., increased dendritic spine density and synaptic protein PSD-95) in both motor cortices, leading to greater M1–M1 functional synchronization (Marchiotto et al., 2025). Thus, in the present study, locomotion was exploited to prompt M1-M1 interhemispheric crosstalk and boost A-tDCS indirect stimulation of the lesioned cortex. This allowed us to directly stimulate an intact cortex instead of a structurally and functionally impaired region (i.e. due to LPC injection), which was modulated indirectly. Notably, although the combined application of A-tDCS and locomotion promoted myelin repair and plasticity, locomotion alone did not induce detectable changes in myelin or oligodendroglia under our experimental conditions. This observation is consistent with our previous work, which showed no plastic change induced by locomotion when this intervention was applied alone (Marchiotto et al., 2025), but clashes with reports indicating that physical activity *per se* can enhance myelin repair (Jensen et al., 2018; Mandolesi et al., 2019; Rossi et al., 2026). Of note, most of these studies employed voluntary running paradigms (i.e., ad libitum access to a running wheel) and intense physical activity (e.g., 5–30 km/day in Rossi et al., 2026), which markedly differ from the standardized, time-limited, and low-/constant-speed locomotion protocol used in the present study (i.e. low-intensity activity).

Importantly, stimulation was applied during a critical post-acute time window corresponding to the late demyelination phase and the initiation of reparative processes. A positive effect on oligodendroglial survival and differentiation was observed as early as 4 days after the stimulation period (i.e. 14 dpi). Stimulated mice exhibited an increase across the entire oligodendroglial lineage, encompassing immature OPCs, premyelinating, and mature OLs in the lesioned cortex. BrdU-based fate-mapping analyses also showed an accelerated loss of immature features and a complementary faster acquisition of markers typical of more advanced stages in oligodendroglial cells. Yet, supernumerary oligodendroglia were not maintained at later time points in the absence of further stimulation, as also occurred to a lesser extent in unstimulated mice. Thus, although stimulated mice tended to exhibit a higher number of OLs compared to unstimulated mice, at 28 dpi, the enhancement of myelin recovery produced by our stimulation protocol was more pronounced than the change in oligodendroglial numbers. It is therefore conceivable that the beneficial effect of A-tDCS+walk involved the production of a higher number of myelin internodes and/or of longer internodes, similar to what has been observed following low intensity rTMS in the mouse motor cortex after cuprizone-induced demyelination (Nguyen et al., 2024). Likewise, increased myelination in the superficial layers of the directly-stimulated uninjured cortex was not accompanied by any increase of immature or mature oligodendroglia, pointing to *de novo* myelin formation and/or remodeling of existing internodes as possible substrates of the stimulation-induced effects.

Several mechanisms may underlie these effects. In the directly-stimulated cortex, A-tDCS might exert a direct action on oligodendroglia, which express a repertoire of voltage-gated ion channels and receive synaptic input from neurons (Butt et al., 2025), thus being exceptionally equipped - among glial cells - to sense and respond to changes in their local electrical environment. In line with this and with OPC hypertrophic arborization detected in the directly-stimulated cortex (see Fig. 6D,E), a recent study has shown that OPC processes preferentially contact nearby cFos+ active neurons in the mouse hippocampus, suggesting that neuronal activity is a key determinant for OPC process targeting and arborization (Sun et al., 2025). Moreover, in both injured and directly-stimulated cortex, A-tDCS may act indirectly on oligodendroglia by increasing neuronal activity and driving the release of activity-dependent signals, such as glutamate, GABA, nitric oxide or BDNF (Barbati et al., 2020; Fritsch et al., 2010; Heimrath et al., 2020; Podda et al., 2016; Rossi et al., 2026; Zhao et al., 2020), which are known to regulate oligodendroglia survival and maturation, and could also account for the elaboration of longer/more abundant internodes (Fletcher et al., 2018; Garthwaite et al., 2015; Liñares et al., 2006; Maas and Angulo, 2021). Glial cell interactions should also be considered, especially in the context of myelin injury, where stimulation could modulate microglial and astrocytic responses, creating a more permissive environment for repair (e.g. enhancing myelin debris clearance or influencing the release of pro-inflammatory cytokines; (Cherchi et al., 2022; Gellner et al., 2021; Monai et al., 2016; Rossi et al., 2026, 2024; Williams et al., 2022).

The observed effects of stimulation on myelin repair and plasticity in the mouse motor cortex reveal a previously unrecognized biological substrate for the therapeutic and neuromodulatory actions of tDCS, with important functional implications. The primary motor cortex M1 produces precisely coordinated neuronal activity patterns essential for skilled motor behavior (Guo et al., 2015) and is one of the most highly myelinated regions of the cerebral cortex (Glasser and Essen, 2011; Redlich and Lim, 2019). Former studies have shown that myelin loss perturbs neuronal activity and synchrony in the mouse motor cortex, resulting in significant motor dysfunction (Bacmeister et al., 2020; Gagnon et al., 2025), whereas activity-dependent myelination is an active regulator of motor circuit function, being critical for the consolidation of learned movements (Bacmeister et al., 2022; McKenzie et al., 2014). Among cortical neurons, parvalbumin-expressing interneurons are remarkably dependent on myelin to support high-frequency firing (Dubey et al., 2022), enabling precise inhibitory control that synchronizes principal excitatory neurons and shapes cortical oscillatory dynamics (Estebanez et al., 2017; Gagnon et al., 2025). In humans, myelination metrics have been reported to correlate with action switching, likely *via* modulation of local primary motor cortex dynamics (Lazari et al., 2022). Thus, by enhancing remyelination and modulating internodal architecture - especially in supragranular layers, which contain a mixed population of intracortical and projection axons from local interneurons, ipsi-/contra-lateral layer II/III pyramidal neurons and thalamus (Chovsepian et al., 2017; Hooks et al., 2013; Marchiotto et al., 2025; Tremblay et al., 2016) - tDCS may increase action potential conduction velocity and fidelity, thereby influencing synchronization of neural networks. In line with this, in our study, stimulation resulted in increased M1-M1 synchrony across all frequencies, but especially in the low-theta (4-8 Hz) and slow-gamma (20-30 Hz) frequency ranges. Of note, in rodents, low-theta frequency is a hallmark of motor network activity, especially linked to locomotion (Noga et al., 2017; Vanni et al., 2017), whose preservation likely reflects better motor functions upon myelin loss.

Finally, our findings place oligodendroglia/myelin plasticity as part of tDCS-induced structural changes, which, so far, have been mostly characterized at the level of synapses and dendritic spines (Barbati et al., 2022; Marchiotto et al., 2025). Whether these distinct forms of structural plasticity merely occur in parallel or are causally linked remains unclear. Evidence in favor of the second hypothesis has been recently provided by Pepper and colleagues (2025), who showed that preventing adult myelination (i.e. generation of new mature OLs) alters the ratio between plastic thin *vs.* stable mushroom spines along the basal dendrites of pyramidal neurons in the mouse motor cortex. Interestingly, in our study, increased myelination was detected only in the supragranular layers of the uninjured directly-stimulated cortex, where increased numbers of spines along the apical and basal dendrites of pyramidal neurons have been formerly found in a similar stimulation paradigm (Marchiotto et al., 2025). As the intensity of the tDCS electric field decreases exponentially with depth from the cortical surface (Rush and Driscoll, 1968), the co-occurrence of these two plastic phenomena in the uppermost cortical layers might be explained by their proximity to the electrode. Yet, accumulating evidence that oligodendroglia can directly shape circuit architecture in an activity-dependent manner through axonal remodeling and synapse engulfment (Buchanan et al., 2023) strongly suggests that a direct functional link between dendritic spine plasticity and oligodendroglial/myelin dynamics warrants further investigation.

In conclusion, although caution is required when extrapolating findings from mice to humans, our data support the potential of A-tDCS as a therapeutic strategy for conditions associated with myelin damage. These findings prompt further studies aimed at elucidating the mechanistic basis of tDCS-induced effects and optimizing stimulation parameters (e.g., intensity, duration, number of sessions, time window) to guide its clinical application.

## Acknowledgements

We wish to thank Dr. Roberta Parolisi (Dept. of Neuroscience Rita Levi Montalcini) for the precious assistance in confocal microscopy analyses. This study was supported by Ministero dell’Istruzione, dell’Universita e della Ricerca—MIUR (Italy) project “Dipartimenti di Eccellenza 2023–2027” to Department of Neuroscience “Rita Levi Montalcini of the University of Turin. MC was partially supported by FISM - Fondazione Italiana Sclerosi Multipla - cod. 2020/PR-Single/028 and financed or cofinanced with the ‘5 per mille’ public founding and by the Next Generation EU/Ministry of University and Research project: “A multiscale integrated approach to the study of the nervous system in health and disease (MNESYS)”, CUP B33C22001060002, PE00000006 missione 4, componente 2, investimento 1.3.

